# Exhaustive reconstruction of the CRISPR locus in *Mycobacterium tuberculosis* complex using short reads

**DOI:** 10.1101/844746

**Authors:** Christophe Guyeux, Christophe Sola, Guislaine Refrégier

## Abstract

Spoligotyping, a graphical partial display of the CRISPR locus that can be produced *in vitro* or *in silico*, is an important tool for analyzing the diversity of given *Mycobacterium tuberculosis* complex (MTC) isolates. As other CRISPR loci, this locus is made up of an alternation between direct repeats and spacers, and flanked by *cas* genes. Unveiling the genetic mechanisms of its evolution requires to have a fairly large amount of fully reconstructed loci among all MTC lineages.

In this article, we point out and resolve the problem of CRISPR reconstruction based on short read sequences. We first show that more than 1/3 of the currently assembled genomes available for this complex contain a CRISPR locus erroneously reconstructed, and errors can be very significant. Second, we present a new computational method allowing this locus to be reconstructed extensively and reliably *in silico* using short read sequencing runs. Third, using this method, we describe new structural characteristics of CRISPR locus by lineages. We show how both the classical experimental *in vitro* approach and the basic *in silico* spoligotyping provided by existing analytic tools miss a whole diversity of this locus in MTC, by not capturing duplications, spacer and direct repeats variants, and IS*6110* insertion locations. This description is extended in a second article that presents general rules for the evolution of the CRISPR locus in MTC.

This work opens new perspectives for a larger exploration of CRISPR loci diversity and of mechanisms involved in its evolution and its functionality.

## 1. Introduction

The CRISPR locus of *Mycobacterium tuberculosis* complex (MTC) was first described in 1993 under the “Direct Repeat” locus designation [1]. It is made of 36 nucleotides-repeats interspaced by unique spacers of a mean of 37 bp (interval: 37bp-45bp). The repeats were soon themselves designated as Direct Repeats and abbreviated as such (DR). The two first sequenced isolates gave access to 43 different spacers sequences. The detection of their presence/absence soon led to the development of the innovative “spoligotyping” method [2]. This method became very popular by its ease and digital format, and was indeed instrumental to decipher the global population structure of MTC [3]. More recently, Whole Genome Sequencing (WGS) studies confirmed that for many lineages and sublineages, the spoligotyping signature allows a correct taxonomical assignment [4], although some generic signatures remains either meaningless, imprecise or convergent, thus largely justifying the use of SNPs as preferred taxonomical markers either globally [5], or for Lineage 4 [6], for Lineage 1 [7], or for Lineage 2 [8].

As in other species with functional CRISPRs, this locus is accompanied by a set of CRISPR associated (*cas*) genes. Their number and nature makes MTC CRISPR type fall into Type III-A group inside CRISPR-Cas taxonomy. It was recently shown to be active under its H37Rv form [9, 10]. Yet, part or the entire region is deleted in several MTC sublineages [11]. Whether the deletion of some of the *cas* genes in the CRISPR-Cas locus may promote genomic instability in some epidemic strains of MTC is another important issue [12].

The genomic diversity of the CRISPR locus has been investigated in detail in a previous study on genomic diversity suggesting that spacer duplication, spacer variation, and IS*6110* insertion sites could be found in the various phylogenetical lineages of MTC [13]. However, it concerned a collection of only 34 MTC strains and did not include any investigation on *cas* genes. Understanding evolutionary dynamics of this locus requires exploration of CRISPR-Cas region on an extended dataset. While the classic *in vitro* approach to spoligotyping is very easy to perform on large datasets, it only provides information on the presence or absence of a well-known list of spacers. This approach does not allow us to know for instance if the order of the spacers may change from one strain to another. Neither does it reveal if there has been a duplication of part of the locus. Finally, it does not provide information on the presence of insertions such as IS*6110*, nor on the existence of single nucleotide polymorphism (SNP) in its direct repeats or spacers. This masks potential functionally significant changes in the loci, and makes it impossible to carry out thorough evolutionary studies. New *in silico*-*based* approaches were developed to produce spoligotypes based on genome reads (SpolPred, spoTyping), however these methods similarly focused on the presence/absence detection of the spacers, so that they have the same limitations as those pointed out above for *in vitro* method [14, 15].

The increased availability of whole genome sequencing of MTC is very promising, insofar as the reads covering multiple times all the places of the genomes, contain these SNPs, duplications, and possible insertions of the CRISPR locus. While, on the whole, it is easy to reconstruct most of a tuberculosis genome using traditional read assembly tools such as Velvet [16], this reconstruction is much more difficult for the CRISPR locus. Indeed, in this part of the genome, the same DR sequence is found between each pair of spacers. Since the size of a DR is not far from that of the k-mers usually used during assembly, there is a risk of wrong bifurcations when searching for a Eulerian path in the De Bruijn graph associated with this assembly. In this context, duplications are difficult to detect, especially when IS*6110* insertions in the locus increase the risk of errors.

In this article, we present a new method to reconstruct CRISPR-Cas systems of MTC from raw Illumina (Illumina Inc, Sand Diego, CA) sequencing runs under a semi-automatized process. It is reliable and robust provided that the reads have sufficient coverage and sizes long enough to span more than one DR. This tool, based on the analyses carried out in [17, 18] particularizes the De Bruijn approach to the specific case of the CRISPR locus and is the main contribution of this article. We show its usefulness both by showing that it can reconstruct CRISPR of reliable reference genomes, and by presenting that mean quality of CRISPR-Cas reconstruction is poor in other assembled genomes available in the public databases. Then we present the high unexpected diversity of the CRISPR-Cas locus of MTC revealed by the exploration of a selection of 434 reads archives. The list of remarkable elements in this locus by MTC lineages is the subject of a separate publication [19].

## 2. Material and methods

### a. Data collection

A first set of data concerns seven reference clinical isolates, for which both assembled genomes and short reads sequencing runs were available, downloaded from the NCBI website, and renown as reference strains (**Table 1**). This selection was made with the a priori that assembled genomes would be highly reliable. This concerns the following strains: CDC1551, Erdman, F11, H37Ra, W-148, and the *M. bovis* BCG str. Pasteur 1173P2 and Tokyo 172.

**Table 1.**
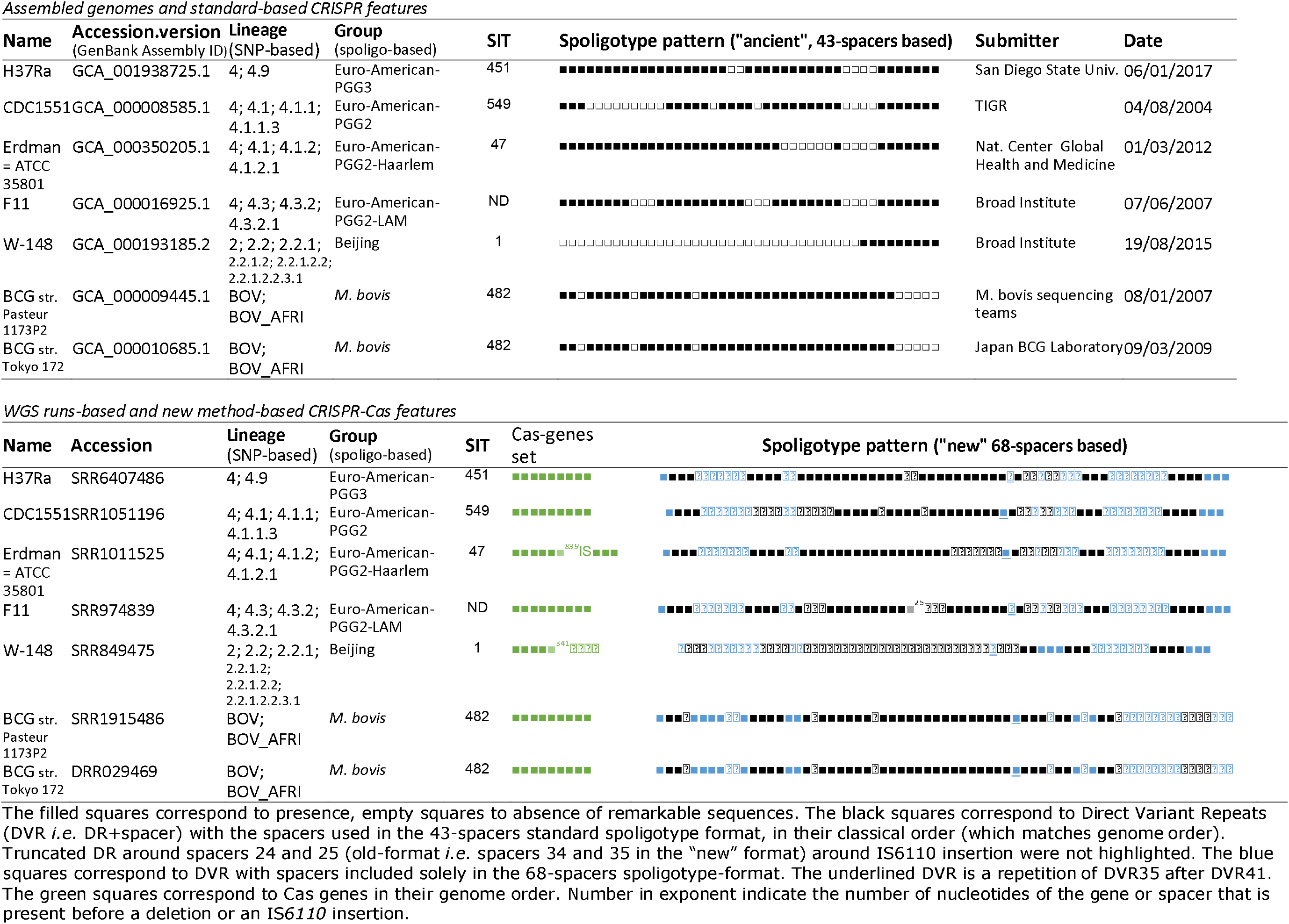
CRISPR-Cas and general features of strains used as references for CRISPR reconstruction pipeline

A second set of data concerns non-reference clinical isolates for which assembled genomes were available but not short reads sequencing runs.

The third set of data comes from a collection of sequence reads archives (NCBI-SRA and EMBL-SRA) that has been retrieved from some state-of-the-art articles to represent the diversity of MTC lineages [20, 21]. This collection was completed by SRA queries on the NCBI search engine, with taxid values of 33894 and 78331, corresponding respectively to *M. tuberculosis variant africanum* and *M. canettii* organisms.

The names of SRA run accessions (SRR) were compiled, then the actual WGS sequencing data were automatically downloaded via the fastq-dump command of the sra-tools package. This led to a database of about 3,500 runs in the form of reads. This database is meant to be a good representative of MTC diversity, both at the lineage level and regarding geographical origins.

A first selection on these runs was carried out, first of all concerning the sequencing technology, which should have been paired-end Illumina to avoid having to manage different formats in our scripts. We also recovered the size of the reads and the average coverage, and discarded all runs corresponding to weak covers (<50×) or with reads too small (minimum size of reads: 75 bp). This collection, once cleaned, was automatically annotated using the script described below, in order to attribute to each run its lineage, its spoligotypes “old format” (43 spacers) and “new format” (98 spacers), as well as its Spoligo-International-Type (SIT) as described in [22].

### b. Runs annotation

As a first annotation of the short sequencing runs (WGS data), we assigned the lineage/sublineage, for each single nucleotide polymorphism (SNP) referenced in [23] for all lineages, in [6] for L4 sublineages, in [8] for L2 sublineages, and in [7] for L1 sublineages. The annotation was made automatic by a script written in Python language that extracts, from its position in the reference genome H37Rv, a neighbourhood of 41 nucleotides centered around each SNP. For each run and each lineage-defining SNP, this 41 base pair sequence was then blasted on the read sequences (blastn, maximum e-value 1e−5, from a local blast database calculated for each genome). At each blast output, we then counted the number of matches that contain the 41 bp version of H37Rv, and the number of matches that contain this pattern whose central nucleotide has been replaced by the SNP tabulated in [5–8]. If the number of mutated units was significantly higher than that of reference H37Rv, the line associated with this SNP was then added to the genome considered.

As a second annotation, we provided the *in silico-derived* old and new formats of spoligotypes based on the presence/absence of known spacers. To this end, we blasted each spacer on each of the read sequences (blastn, e-value < 1e−6), and we calculated the number of matches for each spacer (without looking at whether the sequences matched exactly, as spacers could have been mutated): if this number of matches exceeded 5% of the mean genome coverage, then we considered that the spacer could be added to the spoligotypes. At this level, the percentage has been preferred to a simple occurrence, because, for a certain number of runs, some spacers appeared in 2 or 3 reads when the number of occurrences of the other spacers exceeded, e.g., 70 – and this phenomenon tended to increase with coverage. These few spacers must obviously correspond either to a contamination, to a minor strain in a double infection, or less likely to similarities that appeared by chance due to reading errors, the latter increasing with the number of reads. As for the threshold value for the percentage, it was set in this way after various tests, and by comparing the spoligotypes produced with those known for reference strains. The SIT could then be deduced from a correspondence table derived from SITVIT2 [22].

### c. Assembled genomes annotation and analysis

Slightly adapted script from what was set-up for runs were written to annotate these genomes in term of spoligotype profiles and lineage/sublineage assignation. MIRU-Profiler was used to infer MIRU types from assembled genomes [24]. Resulting patterns were entered in TBminer to identify most likely MTC sublineage assignation according to MIRU-VNTR or spoligo profile or their combination [25].

### d. Listing of CRISPR-Cas remarkable sequences

#### i. Direct repeats and spacerss

In order to evaluate the presence of direct repeats (DRs) and spacers variants, we first needed the list of the reference ones. We thus compiled a first catalogue of the corresponding reference sequences that will be later inflated with variants. DR0 is the name given for reference DR [13], reference spacers k are referred to as esp_k_.

We then looked for spacer variants, using regular expressions in randomly picked up runs from the sample #3 database. More specifically we searched in all the reads for patterns made up of : the last 12 nucleotides of the DR0 [13], followed by a variable sequence with a size between 10 to 70 nucleotides, followed by the first 12 nucleotides of the DR0 (findall method of the python re module, patterns: DR0[−2:]([ATCG]{10,70})[DR0:12] and its reverse complement). The subsequences thus produced were then compared to the reference spacers as soon as they exceeded a number of occurrences fixed according to the coverage: if a given subpattern frequently appears between these two sections of DR0, and if it is not part of the known spacers, then it is determined whether it is a new spacer or a variant of a known spacer, in the following manner. The known spacer most similar to the detected subpattern is looked for, using a Needleman-Wunsch editing distance (compatible with substitution, indels, and gap insertion operations). If this similarity is greater than 95%, the subpattern is considered to be a variant of the most similar spacer and is integrated as such in the catalog with a label of the following type esp_k_(*i*) where *i* is the variant rank; otherwise, it is integrated in the catalog as a new spacer as esp_l_ where l=previous spacer number +1, see algorithm #1 in **Supplementary file 1**.

We then use this enhanced catalogue of spacer variants to find DR variants, in the same way as above. For each pair of spacers esp_k_, esp_l_, for k,l = 1…98, we look in the reads for subunits consisting of the last 12 nucleotides of esp_k_, followed by 30 to 40 nucleotides, themselves followed by the first 12 nucleotides of esp_l_. Again, reverse complement was considered, to double the number of matches, and the possibility of a “\n” for reads spread over more than one line was also included. The new DRs thus obtained were then used in a second phase of discovery of spacer variants, as before, taking into account that the sequences bordering on spacers can be variants of the DR0.

#### ii. Other sequences of interest

To this collection of subpatterns of interest to be discovered in the CRISPR loci, we added:

1. the beginning and end sequences of IS*6110* and its reverse complement (40 bp each time).
2. CRISPR approximate borders: sequences corresponding to Rv2816c (*cas2* gene of the cas locus) and Rv2813c, reputed to border the CRISPR locus [10].
3. CRISPR exact-flanking sequences: the reads including the end of the *cas2* gene have been extracted from a small collection of genomes from the database presenting spacer 1 in its “new spoligotype” to retrieve likely ancestral closest border to CRISPR locus. The consensus sequence located downstream has been reconstructed, then the reads including the latter were recovered. These included a DR0 followed by the spacer “new” number 1. After verification (blast), this CRISPR-flanking pattern was indeed found in a large set of genomes in our collection, so it was added as such to the catalog of patterns of interest. The same treatment was performed on genomes with spacer 68 to identify the end sequence between the latter spacer found in MTC stricto sensu (without *M. canettii*) and the Rv2813c gene. The corresponding pattern was also added to our catalog.

### e. Locus reconstruction

#### i. Contiguage

For each run, the sequences of interest mentioned above were first blasted on all the reads (blastn, evaluated 1e−7), in order to extract the small set of reads potentially covering the CRISPR locus. This small set of reads was then extended, where each read of size n was transformed into its n−k+1 k-mers, where k is equal to the integer part of 4n/5. This step, inspired by a classical contiguage by De Bruijn approach [26], was carried out on the one hand to have a good coverage of CRISPR in terms of k-mers, and this even if the original coverage was close to 50×, and on the other hand, in order not to definitively disqualify for the next steps a read with a possible reading error: in what follows, only its k-mers containing this error will be disqualified. Corresponding algorithm is available in **Supplementary file 1** (algorithm #2).

A sequence is thus randomly extracted from this set of k-mers potentially covering the CRISPR, serving as a starting point for the first contig, to which an initial score of 1 is associated. The k-mers such that their first k-1 nucleotides correspond exactly to the last k−1 nucleotides of the current contig are then obtained from the set of k-mers. It is then regarded if the majority of the latter have the same last nucleotide (i.e., in position k). If this is the case, this nucleotide is added to the current contig, the k-mers that have matched are removed, their number is added to the score of the current contig, and the progress continues to be made in the reconstruction of the locus with the next nucleotide. If this is not the case, we start again with the other side of the current contig, looking for k-mers whose last k−1 nucleotides correspond exactly to the first k−1 nucleotides of the current contiguous. And the latter is no longer extended from his tail, but from his head.

At a time when no consensus seems to be emerging for the new nucleotide to be added to the current contig, this latter is stored separately with its score, and the whole process is repeated from a new randomly extracted k-mer. As, at each iteration, at least one k-mer is removed from the original set, this process has an end, leading to a more or less long list of potential contigs, themselves more or less long.

The contigs are then manually processed by decreasing score, in order to reconstitute the CRISPR structure. To this end, the catalogue of sequences of interest (variants of spacers and DRs, sequences bordering the IS*6110*, and the start and end patterns of the locus) is iterated, in order to replace each nucleotide sub-sequence by its name using the replace method of the str class (python). The result of this post-processing of the previously obtained contigs is a reasonably sized character string, including patterns of the form *spX(Y)* for the variant Y of the spacer X, *DRX* for variant number X of the DR, as well as the words *begin_IS6110*, *end_IS6110*, *begin_IS6110c*, *end_IS6110c*, *starting_pattern*, *ending_pattern*, *Rv2816c*, and *Rv2813c*. This translation makes it easier to understand the contigs obtained, and makes it easy to detect a break in the order in which the spacers appear. It also allows to detect new variants that had not been detected until now, and to add them after naming to the database of remarkable sequences. In the vast majority of the cases studied manually (but read exceptions in Duplication paragraph below), one to three contigs depending on the number of IS6110 insertions in the locus (those with the highest scores) were sufficient to reconstruct the entire locus. The extreme elements of said contigs always were either the sequences bordering the locus or a beginning or end of IS*6110*(c).

#### ii. Duplications resolution

If the reconstruction, mentioned above, of the CRISPR locus makes it possible to highlight the tandem duplications of spacers, in the case of read files of size >75 (leading to k-mers >56 bp, as in our selected WGS data), it nevertheless passes through possible duplications spread over several spacers. Let’s suppose that we have a pattern of the form: sp_k_*sp_k+1_*…*sp_l_*sp_k+1_*…*sp_l_*sp_m_. Then, once the contig is rebuilt to the end of spacer number l (and its DR), what comes next in the reads concerns both sp_k+1_ and sp_m_: when these two sequences diverge, there is no longer a nucleotide consensus in the considered reads, and the expansion of the contig stops. In addition, the k-mers of the second repeated pattern were used in the expansion of this contig when it was at the first pattern, to a number of k-mers used and removed twice as large as expected, and to the impossibility of reading the repetition of the pattern.

At this stage, we can conclude that if the expansion of a contig has not stopped on an IS or a sequence bordering a CRISPR locus, and if the score of said contig is higher than expected, then there is a suspicion of large-scale duplication. To resolve this situation, post-treatment was added to the locus reconstruction pipeline: for each pair of spacers (k,l), k,l=1…98, we count the number of k-mers containing the last 12 sp_k_ nucleotides, followed by any of the DR variants, followed by the first 12 sp_l_ nucleotides. And couples whose number of matches is significant are displayed in lexicographic order. In this list, a pattern of the form spl*DRX*sp_m_, l≥m, is proof of a duplication (in tandem when l=m): after l, we loop back to m<l. Of note, the successions of spacers involved in this duplication have a number of k-mers of the order of twice the successions of spacers located outside this duplication. And this doubling of the number of matches is a form of cross-validation of the duplication.

At this stage, we are therefore able to reconstruct the entire CRISPR locus from Illumina paired-end reads, provided that the coverage and size of the reads are reasonable, and this by being able to detect duplications, spacer and DR variants, and IS insertions. This process is 95% automated, but it requires human intervention to finalize the assembly of the contigs. Once this locus has been reconstructed, the resulting spoligotypes (old and new) can be compared to spoligotypes based on presence/absence of spacer sequences. The algorithm is shown in **Supplementary File 2**.

### f. Runs’ additional selection

A final point remains to be clarified at this stage, namely how the WGS runs here reconstructed were selected from our database of ~3,500 items. Indeed, although much of the reconstruction has been automated, the remaining 5% takes a little time to be properly carried out. Not wanting to waste time rebuilding loci where nothing has happened, in terms of insertion and duplication, we have taken part of the pipeline detailed above to make a selection of the runs of interest. These correspond to samples carrying duplications as well as samples carrying IS*6110* insertions.

For a given run, we focus on reads returning matches during a blast on sequences of interest (DR and spacers). This again is performed using k-mers derived from the reads as described above. Then, patterns of the shape of an end of spacer l, followed by a variant of DR, itself followed by a beginning of spacer m, where l≥m, are looked for, as they are signs of duplication. Similarly, patterns of the form end of spacer k, followed by 0 to 36 nucleotides, themselves followed by the beginning of IS*6110*, are looked for insertions in DRs. Finally, ends of DR variant, followed by a certain number of nucleotides, and then the beginning of IS*6110* for insertions, are searched for insertions in spacers (with all possible variations in terms of layout and reverse complement). Only runs with either of these conditions were further considered, as basis of knowledge for the numerical study detailed below.

## 3. Results

### a. Evaluation of CRISPR locus reconstruction based on WGS data of MTC reference strains

We first reconstructed the CRISPR loci of the best MTC studied strains using corresponding sequencing runs. Although it should be noted that these 7 reference strains do not represent the full MTC diversity since only four lineage 4 strains, two *M. bovis* BCG variants, and a single lineage 2 strains are concerned (**Table 1**). Still they concern three distant lineages among of the 7 lineages constituting MTC diversity.

Briefly, we blasted the subsequences that are part of CRISPR-Cas locus (referred to as “remarkable sequences”) against the sequence reads against. These reads were then used to build contigs by the De Bruijn approach [26]. During contig building, scores were calculated taking into account the number of reads involved. Contigs included exclusively remarkable sequences so that their structure could be coded as the list of the corresponding tags. Note that numbering of spacers are by default those from the 68-spacers format referred to as “new format” in this article [13]. The contigs were then processed manually in decreasing order of scores to resolve possible duplications and sequences flanking IS*6110* insertions. The CRISPR structure was then coded as a binary pattern listing the presence or absence of the remarkable sequences in their order of appearance (spoligo-like profile) (**Table 1**, **lower part**).

For assembled genomes, we first identified the location of CRISPR locus using one of the remarkable sequences. The whole locus was then extracted and translated both as the list of actually present remarkable sequences, and as a binary pattern in a spoligotype-like format. The classical 43-spacers spoligotype was then extracted considering only the useful information (Table 1 **upper part**).

With both WGS-derived and assembled genomes-derived CRISPR features, we found the same spacer sequences alternating with the same DR sequences. This was true for DR variants found between spacers 25 and 26 (truncated version), between spacers 30 and 31, between spacers 64 and 65, 66 and 67, and between spacers 67 and 68 as described previously [13]. We also identified the expected IS*6110* sequence in the DR between spacers 34 and 35. Last, we detected a duplication of spacer 35 and the adjacent DR (Direct Variant Repeat 35 or DVR35) as described by van Embden *et al*, but we always identified it at the 3’ end of DVR41, not DVR45 as described in text by these authors [13].

At the level of the spacer variants, a single discrepancy was identified around spacer 13 in H37Ra: in the assembled genome, there is a variant of the spacer with 10 more nucleotides, corresponding to tandem duplications of nucleotides, itself surrounded by two distinct variants of DR, one with a size 46 and the other with a size 39. These supposed DR inflations again correspond to tandem nucleotide duplications.

Altogether, the CRISPR-Cas locus reconstructed by our pipeline using WGS of reference strains matches perfectly with the public assemblies. This validates our analytic pipeline to annotate and reconstruct CRISPR-Cas locus based on short-reads runs.

### b. CRISPR region in other MTC assembled genomes

As performed for assembled genomes of reference MTC strains, we extracted the CRISPR-locus from an additional 185 assembled MTC genomes available in the Public databases (sample #2, **Figure 1**). First of all, it should be noted that this sample is far from being representative of the entire *Mycobacterium tuberculosis* complex. Indeed, we find in this set only 8 genomes of L1, two of L3, and neither L5 nor L7. Moreover, among the well represented lines, the diversity in terms of sublineages is not respected at all: we find only sublineage 2.2 in the 44 genomes of lineage 2 (including 40 of 2.2.1), when among the 110 genomes of L4, we have 21 of L4.1.2.1, 16 of L4.3.2, 24 of L4.3.3, and for example no 4.6. We also noticed that 25 genomes out of the 187 genomes (~14%) were of really poor quality, accumulating multiple variations of spacers and DRs, at sizes varying greatly. For example, strain GG-77-11, line 4.3.2, has a mutant for spacers 19, 20, 21, 25, 32, 34 and 42. Other genomes with high frequency in spacer variants were EAI5_NITR206, CAS_NITR204. In these assembled genomes, we also occasionally found spacers 46 and 48 under various forms (variants) and places. We also noticed that of the 27 assembled genomes of 4.1 or 4.2 with the pair of spacers 41 and 42, 24 genomes failed to duplicate the 35 after the spacer 41. At the IS*6110* level, all assembled genomes have an insertion upstream of spacer 35. However, only one other IS*6110* copy was identified, in front of spacer 46 of strains of sublineage L2.2.1.

**Figure 1:**
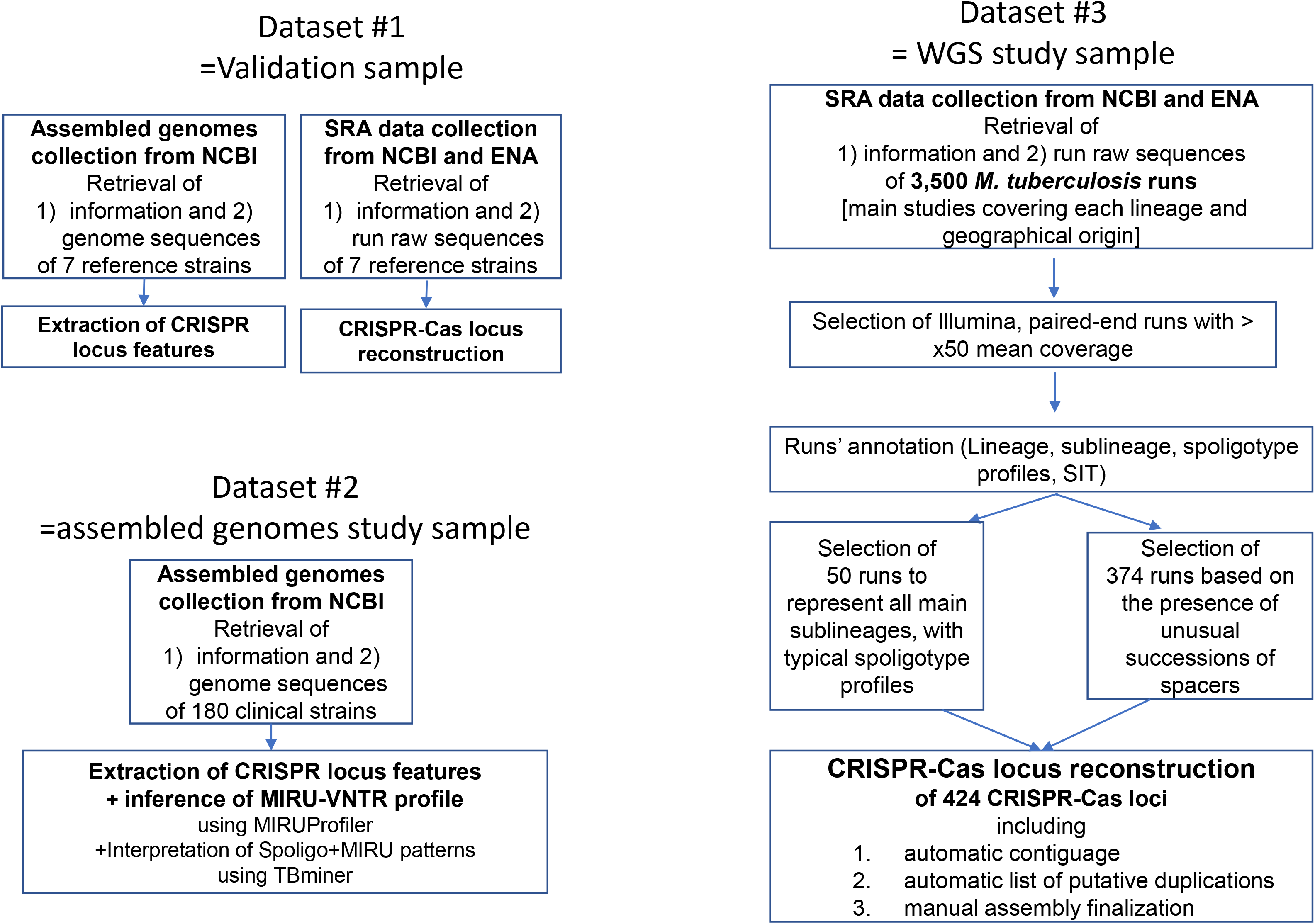
Diagram showing the different dataset of our study and the process of MTC CRISPR Locus reconstruction

We then derived their 43-spacers spoligotype patterns. This profile was interpreted in terms of classification using TBminer, and robustness of this classification was further explored using MIRU-VNTR patterns derived from MIRU-Profiler (**supplementary file 3**). All samples had sufficient information to enable their classification according to spoligotype patterns, and this classification was found convergent with MIRU-VNTR patterns for almost all of them (n=174, 94%). In parallel, we used our annotation procedure to classify all samples according to SNPs. Most of them exhibited several SNPs, and almost all sublineages SNPs confirmed lineage classification (samples carrying L1 SNPs carried L1-sublineages SNPs and not SNPs from sublineages from, let’s say, L4 lineage). We then compared spoligotype-derived classification to SNPs-derived classification. Surprisingly, among the 65 non-L4 samples according to SNPs, 13 samples (belonging either to L1, L2 or L3) were classified as L4 according to their spoligotype and MIRU-VNTR patterns (**Table 2**). They indeed presented the typical sp43-50 (sp33-36 in the ancient format numbering) deletion characteristic of the L4 lineage that, until now, has never been described for strains of other lineages to our knowledge.

**Table 2.**
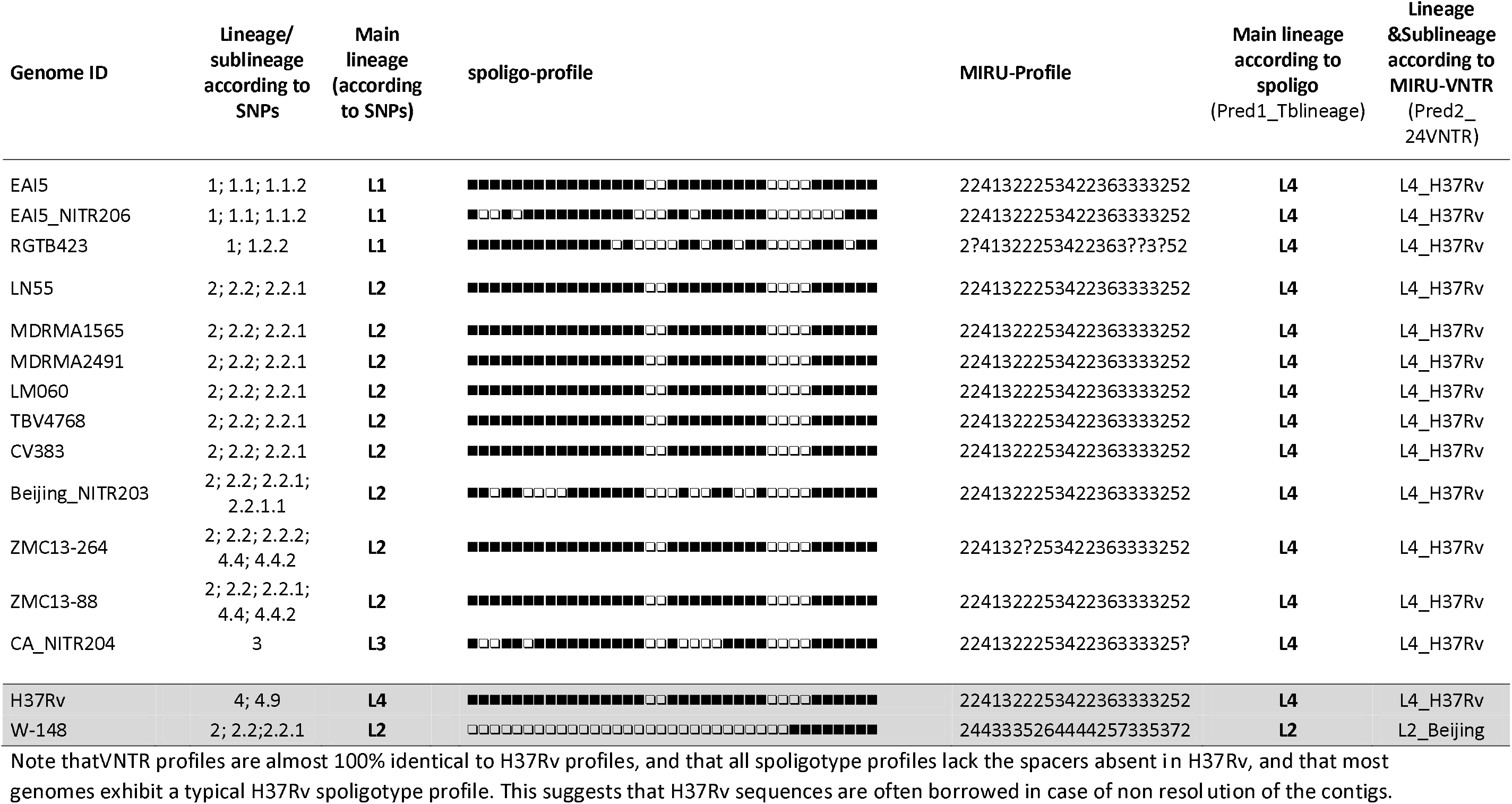
Spoligotype and MIRU-VNTR profiles of Public assembled genomes with discrepancies between SNP-based and spoligo-derived classification

In addition, among the 112 L4 samples, 54 samples had a typical MIRU-VNTR and spoligotype profile characteristic of H37Rv without belonging to L4.9, the H37Rv specific sublineage according to SNPs. They indeed carried the typical del sp30-31 (20-21 in ancient format) deletion characteristic of the H37Rv sublineage inside L4.9 (**Supplementary file 3**). Altogether we identified 67 assembled genomes (36%) with clear discrepancies between CRISPR and MIRU-VNTR information and SNPs, with many instances where reference genome sequences seem to have been borrowed and included in the assembly: a wide proportion of assembled genomes have likely erroneous CRISPR-loci, impeding their exploration to understand CRISPR diversity and evolutionary dynamics.

### c. CRISPR evolutionary events in MTC

We reconstructed the CRISPR-Cas locus of 434 strains representing the diversity of the MTC lineages and showing interesting features (**Figure 1**, sample #3). The global CRISPR profiles obtained were found to be consistent with SNPs-derived lineages and sublineages (del 43-50 found in L4 samples, etc., **Table 3**, **Supplementary file 4**). The resulting data are a pre-requisite to infer general principles of evolution in this part of the genome. As explained previously, these results and lessons will be the subject of a companion article [19]. In what follows, we will use these teachings to compare our method to the prevailing Velvet-based one [16]. To this end, we list the different types of events made detectable by the aforementioned method. They have been systematically observed in all lineages, in one or more lineages, or in a clearly defined sub-lineage.

**Table 3.**
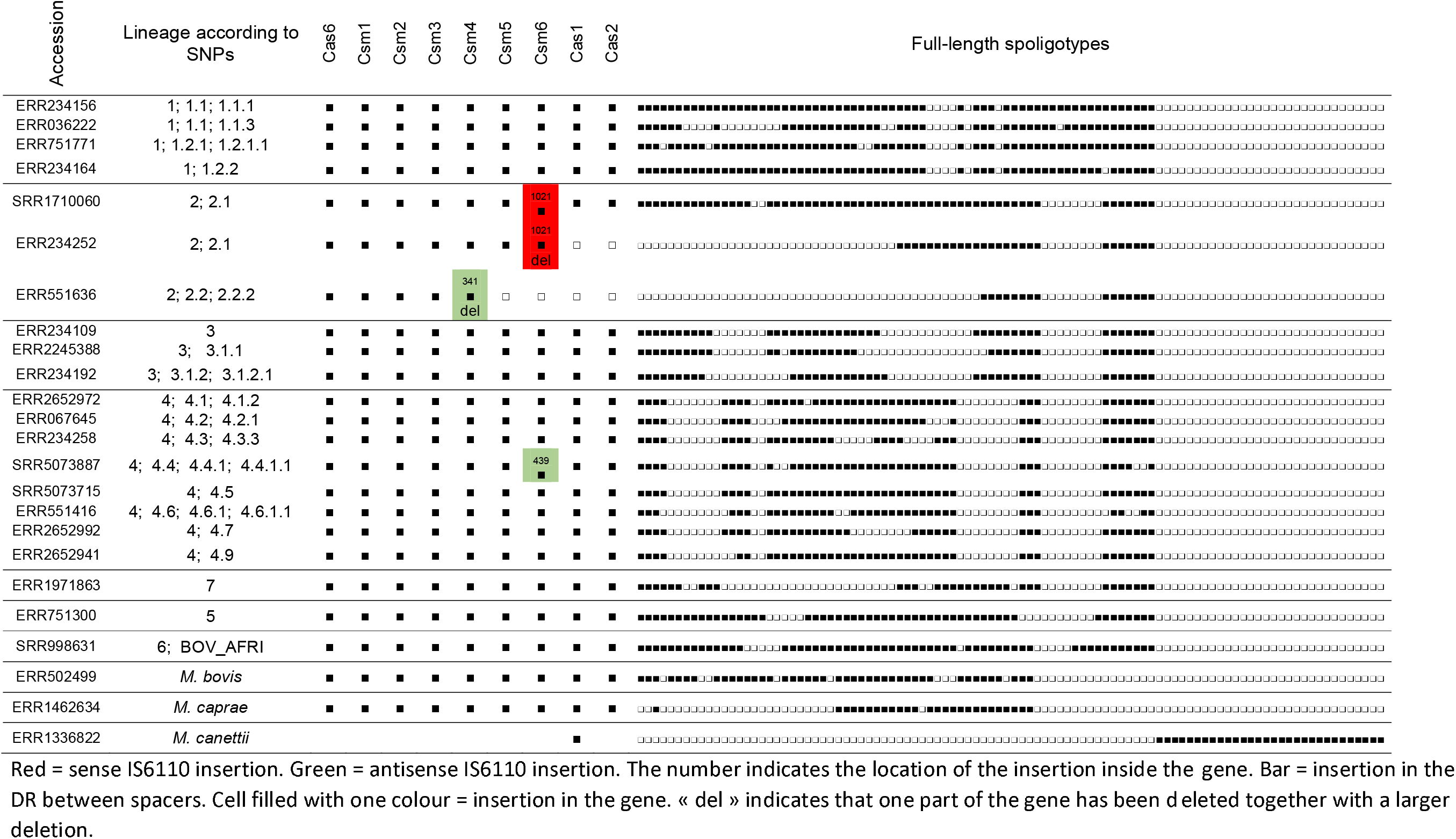
CRISPR-Cas locus profile reconstructed from pubic WGS runs and representative of MTC diversity

Regarding DR diversity, almost all the time, there is the same direct repeats (DR) sequence between two given spacers. The DR0 version of the DR is largely predominant. The exceptions observed in the 7 references strains were confirmed:

- Regardless of the strain, the same variation between spacers 30 and 31 is always found (DR2). A second variant is systematically found between spacers 66 and 67 (DR4), and a third between spacers 67 and 68 (DR5), and a fourth one between sp64 and 65 (DR6, **Table 4**).
- All L1 samples, and only they, have an original DR variant between spacers 50 and 51 (DR3), and those of sublineage L1.1.1.1 have another variation between spacers 14 and 15 (DR1, **Table 4**).
- There are also about 15 other DR mutants, but this is a one-time phenomenon. And if we except the DR between spacers 25 and 26, all DR variants have the same size, *e.* 36 base pairs. The DR truncated between spacers 25 and 26 is identical in all samples that have this pair of spacers.

**Table 4.**
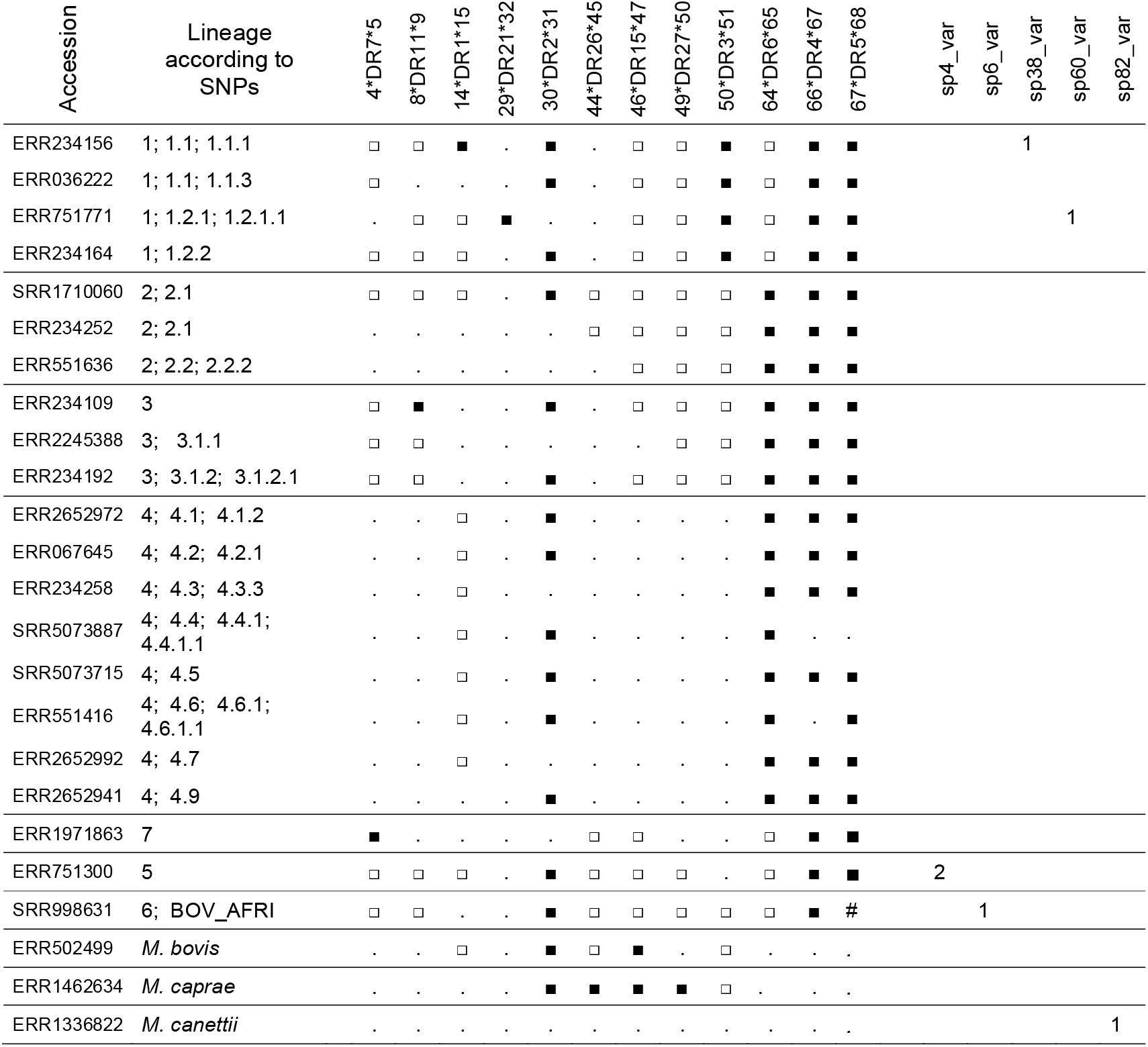
DR and spacer variants for the representative set of MTC diversity

| Accession | Lineage according to SNPs | 4*DR7*5 | 8*DR11*9 | 14*DR1*15 | 29*DR21*32 | 30*DR2*31 | 44*DR26*45 | 46*DR15*47 | 49*DR27*50 | 50*DR3*51 | 64*DR6*65 | 66*DR4*67 | 67*DR5*68 | sp4_var | sp6_var | sp38_var | sp60_var | sp82_var |
| --- | --- | --- | --- | --- | --- | --- | --- | --- | --- | --- | --- | --- | --- | --- | --- | --- | --- | --- |
| ERR234156 | 1; 1.1; 1.1.1 | □ | □ | ■ | . | ■ | . | □ | □ | ■ | □ | ■ | ■ |  |  | 1 |  |  |
| ERR036222 | 1; 1.1; 1.1.3 | □ | . | . | . | ■ | . | □ | □ | ■ | □ | ■ | ■ |  |  |  |  |  |
| ERR751771 | 1; 1.2.1; 1.2.1.1 | . | □ | □ | ■ | . | . | □ | □ | ■ | □ | ■ | ■ |  |  |  | 1 |  |
| ERR234164 | 1; 1.2.2 | □ | □ | □ | . | ■ | . | □ | □ | ■ | □ | ■ | ■ |  |  |  |  |  |
| SRR1710060 | 2; 2.1 | □ | □ | □ | . | ■ | □ | □ | □ | □ | ■ | ■ | ■ |  |  |  |  |  |
| ERR234252 | 2; 2.1 | . | . | . | . | . | □ | □ | □ | □ | ■ | ■ | ■ |  |  |  |  |  |
| ERR551636 | 2; 2.2; 2.2.2 | . | . | . | . | . | . | □ | □ | □ | ■ | ■ | ■ |  |  |  |  |  |
| ERR234109 | 3 | □ | ■ | . | . | ■ | . | □ | □ | □ | ■ | ■ | ■ |  |  |  |  |  |
| ERR2245388 | 3; 3.1.1 | □ | □ | . | . | . | . | . | □ | □ | ■ | ■ | ■ |  |  |  |  |  |
| ERR234192 | 3; 3.1.2; 3.1.2.1 | □ | □ | . | . | ■ | . | □ | □ | □ | ■ | ■ | ■ |  |  |  |  |  |
| ERR2652972 | 4; 4.1; 4.1.2 | . | . | □ | . | ■ | . | . | . | . | ■ | ■ | ■ |  |  |  |  |  |
| ERR067645 | 4; 4.2; 4.2.1 | . | . | □ | . | ■ | . | . | . | . | ■ | ■ | ■ |  |  |  |  |  |
| ERR234258 | 4; 4.3; 4.3.3 | . | . | □ | . | . | . | . | . | . | ■ | ■ | ■ |  |  |  |  |  |
| SRR5073887 | 4; 4.4; 4.4.1; 4.4.1.1 | . | . | □ | . | ■ | . | . | . | . | ■ | . | . |  |  |  |  |  |
| SRR5073715 | 4; 4.5 | . | . | □ | . | ■ | . | . | . | . | ■ | ■ | ■ |  |  |  |  |  |
| ERR551416 | 4; 4.6; 4.6.1; 4.6.1.1 | . | . | □ | . | ■ | . | . | . | . | ■ | . | ■ |  |  |  |  |  |
| ERR2652992 | 4; 4.7 | . | . | □ | . | . | . | . | . | . | ■ | ■ | ■ |  |  |  |  |  |
| ERR2652941 | 4; 4.9 | . | . | . | . | ■ | . | . | . | . | ■ | ■ | ■ |  |  |  |  |  |
| ERR1971863 | 7 | ■ | . | . | . | . | □ | □ | . | . | □ | ■ | ■ |  |  |  |  |  |
| ERR751300 | 5 | □ | □ | □ | . | ■ | □ | □ | □ | . | □ | ■ | ■ | 2 |  |  |  |  |
| SRR998631 | 6; BOV_AFRI | □ | □ | . | . | ■ | □ | □ | □ | □ | □ | ■ | # |  | 1 |  |  |  |
| ERR502499 | *M. bovis* | . | . | □ | . | ■ | □ | ■ | . | □ | . | . | . |  |  |  |  |  |
| ERR1462634 | *M. caprae* | . | . | . | . | ■ | ■ | ■ | ■ | □ | . | . | . |  |  |  |  |  |
| ERR1336822 | *M. canettii* | . | . | . | . | . | . | . | . | . | . | . | . |  |  |  |  | 1 |

At the spacer level, we have the following rules:

- The strains of human and animal L6 lineage (*M. bovis*) all have a mutant of spacer 4, when present.
- The L7 ones all have a variant of spacer 6.
- All strains of lineage 1.1.1.1 have a spacer 38 variant.

Concerning duplications, the following points should be noted: 1) a large duplication between spacers 20 and 21 in lineages 1.1.1.7 and 1.1.1.8; 2) a large tandem duplication of spacer 29 in lineage 1.1.3, as well as spacers 5 and 21 in L3; 3) some of 1.2.1.2 strains have a large duplication of 25 spacers between 57 and 58; 4) there is duplication of spacer 35 everywhere between spacers 41 and 42, with the notable exception of sublineages 4.3 to 4.9 (**Supplementary file 4**).

Finally, as expected, we always found an IS*6110* insertion sequence between spacers 34 and 35. Other IS*6110* insertions were identified in DRs or in spacers, in the sense or antisense direction (**Supplementary file 4**).

## 4. Discussion

We set up a semi-automatic pipeline to reconstruct CRISPR-Cas locus from MTC short reads sequencing runs. We first discuss the robustness of this pipeline and then comment on the problems at stake when trying to reconstruct CRISPR locus using standard assembly pipelines.

### a) Robustness of the pipeline reconstructing CRISPR loci

The pipeline proposed is based on a De Brujn approach and builds contig based on the consensus extension of k-mers. The selectivity of the consensus is cross-validated by the manual exploration of the coverage of the different spacers and DR.

CRISPR loci extracted from MTC reference genomes mainly deriving from Sanger sequencing, and the loci we reconstructed based on short-read sequencing runs of the same samples proved almost 100% concordant. The single discrepancy occurred for H37Ra that exhibited oligonucleotide repetitions in one single spacer and adjacent DR, repetitions that are absent in the highly related H37Rv genome. Two reasons may account for this discrepancy. The first possibility is that the two H37Ra strains actually handled by the two methods were not the same, and rare mutation occurred in the subclone that was used to produce the assembled genome sequence. No such mutation, leading to increased size of a spacer and its adjacent DR, was however observed in the 434 runs explored in the subsequent work, making this possibility quite unlikely. The second possibility is that there was an error during the assembly or the Sanger sequencing used to reconstruct this locus.

The robustness of our pipeline is further supported by the compatibility between SNPs subclustering and clustering derived from specific mutations in the CRISPR-Cas locus, whether they concerned IS*6110* insertions, spacer or DR variations, of duplications. For instance, we observed the sp43-50 deletions in all L4 samples, we observed an IS*6110* insertion downstream of spacer 41 in all 4.1.2.1 samples, a variant in sp4 in all L6 samples, a tandem duplication of DVR5 was observed in all L2 and L3 samples still harbouring this region of the CRISPR (L2.1, most L3) etc. (**Supplementary file 4**). We also could confirm all specificities identified in the pioneer work using targeted Sanger sequencing such as DVR35 duplication for all samples outside L4 and most samples of L4, DR variants between sp30-31, sp 50-51, sp64-65, sp66-67 and 67-68 [13].

This reconstruction results in a high level of additional information as compared to existing methods exploring MTC CRISPR diversity: both *in vitro* and *in silico* spoligotyping only deal with the presence or absence of specific spacers, with methods tolerating non-fully concordant sequences. As a result, all the information of spacer and DR variants, DVR duplications, IS*6110* insertions is lost.

### b) On the use of standard assembly methods for CRISPR reconstruction and resulting assembled genomes in Public databases

The systematic study of the CRISPR loci of the assembled genomes deposited in public databases first showed that, in the approximately 200 genomes currently available, most seem to have a well reconstructed CRISPR, especially the reference samples that likely benefited from partial Sanger sequencing. Conversely, another relatively large proportion of these genomes (more than 1/3) have a clearly problematic locus, not trustworthy at all. This does not mean that there is no benefit in sharing such data, which can be informative for the rest of the genome. However, the problem is that it is difficult to know *a priori* whether, for a given genome, the CRISPR locus is, or is not, trustworthy. The reasons for this average low quality of CRISPR information is first its genetic complexity, and second the difficulty to deal with this complexity when explored using short reads sequences. Obviously, a number of studies have failed in reconstructing this locus using short reads sequencing. The difficulty of such a reconstruction, and the errors that result from it, have their source in several causes, some of which have already been introduced previously.

First of all, the CRISPR locus is by nature a very difficult area to assemble, at least automatically. Indeed, the De Bruijn approaches look for an Eulerian path in the graph whose vertices are the k-mers, and for which there is an edge between two vertices if, and only if a suffix of one is a prefix of the other. This locus contains multiple copies of DRs, IS*6110* insertions, spacers that sometimes share similarities (the beginning of spacer 33 is the end of spacer 36, for example). In addition, we identified common DVR duplications and even large scale duplications, especially in Lineage 1 (Refrégier et al, in preparation). All these events lead to possible bifurcations in the graph.

In addition, the assembly is usually done by Velvet [8], which by default has a maximum k-mer size of 49. In the best case scenario where this size has been set to its preconfigured maximum, knowing that a DR is size 36, this leaves only 13 bp of overlap to be shared between the two spacers, upstream and downstream, which multiplies the incorrect bifurcations in the graph. Increasing this limit value requires recompiling Velvet from its sources, which obviously only a few or no people who submitted their assembled *M. tuberculosis* genomes have done.

Finally, the assembly is often reference-guided. In that case, assembly uses mainly H37Rv, a recent well-studied L4 isolate. However, this strain is not really representative of the diversity of the locus: it has no duplication, and only the ancestral IS copy upstream of spacer 35. When mapping reads to this reference, samples containing spacers not present in H37Rv (such as sp43-50) are likely to be discarded or misplaced. This is why a majority of the spoligotypes derived from the assembled genomes available on the NCBI appeared to be L4-related, while at the SNP level, the lineages were a little bit more diverse: there were obviously holes in the CRISPR locus, which is therefore not trustworthy.

## 5. Conclusion

In this article, we have explained why MTC CRISPR locus should not be assembled using standard tools and we have begun to reveal the unexpected diversity it contains. This was made possible thanks to a semi-automatic method that allows, for genomes with a reasonable coverage and read size, to reconstruct CRISPR-Cas locus in a reliable, fast and robust way. It reveals duplications of various length, variants of spacers and DRs, and insertions of IS*6110* sequences, *i.e.* a full range of evolutionary events that may be found in other CRISPR loci.

In a companion article, we describe the high diversity of MTC CRISPR locus unveiled by our new method, we establish a list of notable elements by lineage, and infer MTC CRISPR various mechanisms of evolution. Among our objectives is the transformation of our tool into a professional quality software, so that the whole community can benefit from it. We also wish to study each lineage separately and in depth, on large sets of representative genomes, in order to reveal the fine evolutionary dynamics of the CRISPR-cas locus.

## Supplementary Material

S1_file : Tables listing the DR, spacer variants and specifically searched patterns in our pipeline, and tables listing SNPs used to infer classification.

S2_file : Algorithm used to reconstruct CRISPR locus (spacer discovery, and contiguage)

S3_file : Spoligotype and MIRU-VNTR patterns of non-reference assembled genomes, derived classification and comparison with classification derived from their SNPs.

S4_file : Selection of reconstructed CRISPRs from short Illumina sequencing runs using our pipeline.

Legend IS6110 sheet. Column A: lineage assignation according to Coll *et al*., Palittapongarnpim *et al.* for L1, Shitikov *et al.* for L2, and Stucki *et al*. for L4; column B to end: the first line designates the name of the gene or the spacer ID; the second title line designates the spacer ID according to the classical spoligotyping nomenclature also visible by yellow color). Vertical bars stand for IS*6110* copies within DR, in green for copies in orientation 1 (antisense) and red for orientation 2 (sense). Color boxes are for insertions within a gene or a spacer, with the same color for the orientation than above. The number in the box indicates the position (nucleotide) where the insertion occurred in the coding sequence. Finally, a large colored tube with white squares depicts a very likely recombination event between two insertion sequences that led to the deletion of all sequences between them (thus, only one IS*6110* remains).

