## Supplementary material for "Exhaustive reconstruction of the CRISPR locus in *Mycobacterium tuberculosis* complex using short reads": Suppl. file 2

```
Supplemental file 2

Algorithm 1: Python-like pseudo-code of new (variant of) spacer discovery

#Set of known spacers
  known_spacers = {i: [sequence(i) for sequence(i) in Van Embden],
               	for i = 1..98}
putative_spacers = []
  for each g in genomes:
  	for each r in g.reads:
      	if s subword of r matches DR0[-12:]([ACGT]{10,70})DR0[:12]:
          	# some read matches the regular expression
          	sp = s[12:-12]
          	if sp not in known_spacers.values():
               	putative_spacers.append(sp)
  considered_sequences = [seq for seq in set(putative_spacers)
                          	if putative_spacers.count(seq) > 50]
  for sp in considered_sequences:
  	if max([similarity(sp, known_spacers[i][0]) for i in 1..98]) < 0.95:
          # A new spacer is discovered
      	    known_spacers[len(known_spacers)+1] = [sp]
  	else:
      	    # id number of the similar spacer
      	    id = idxmax([similarity(sp, known_spacers[i][0]) for i in 1..98])
      	    known_spacers[id].append(sp)


Algorithm 2: Python-like pseudo-code of contiguage
seqs = blastn(sequences_of_interest, reads, evalue = 1e-7)
# k-merization of reads of interest
n = len(seqs[0])
k = int(4*n/5)
seqs = [[seq[u:u+k] for u in 0..n-k] for seq in seqs]
# Contiguage
contigs = []
while len(seqs) > 0:
    contig = choice(seqs)
    seqs.remove(contig)
    forward = True
    while forward:
        # kmers that begin like the end of the contig
        kmers = [seq for seq in seqs if seq[:-1] == contig[-n+1:]
        for kmer in kmers:
            seqs.remove(kmer)
        # A nucleotide is more frequent at the end of the kmers ?
        frequences = {u: [[kmer[-1] for kmer in kmers].count(u) for u in “ACGT”}
        occurrences = sorted(frequences.values())
        if occurrences[0] > 3*occurrences[1]:
            # Adding the most frequent ending nucleotide to the contig end
            new_nucl = idxmax(frequences.values())
            contig += new_nucl
        else:
            forward = False
    while not forward:
        # kmers that end like the beginning of the contig
        kmers = [seq for seq in seqs if seq[1:] == contig[:n-1]]
        for kmer in kmers:
            seqs.remove(kmer)
        # A nucleotide is more frequent at the beginning of the kmers ?
        frequences = {u: [[kmer[0] for kmer in kmers].count(u) for u in “ACGT”}
        occurrences = sorted(frequences.values())
        if occurrences[0] > 3*occurrences[1]:
            # Adding the most frequent beginning nucleotide to the contig head
            new_nucl = idxmax(frequences.values())
            contig = new_nucl+contig
        else:
            contigs.append(contig)
```
